# Choice anticipation as gated accumulation of sensory predictions

**DOI:** 10.1101/2023.12.14.571751

**Authors:** Brandon Caie, Dominik Endres, Aarlenne Z. Khan, Gunnar Blohm

**Affiliations:** Queen’s University; Université de Montréal; Philipps-University Marburg

## Abstract

Predictions are combined with sensory information when making choices. Accumulator models have conceptualized predictions as trial-by-trial updates to a baseline evidence level. These models have been successful in explaining the influence of choice history across-trials, however they do not account for how sensory information is transformed into choice evidence within-trials. Here, we derive a gated accumulator that models the onset of evidence accumulation as a combination of delayed sensory information and a prediction of sensory timing. To test how the delays interact with across-trial predictions, we designed a free choice saccade task where participants directed eye movements to either of two targets that appeared with variable delays and asynchronies. Despite instructions not to anticipate, participants responded prior to target onset on some trials. We reasoned that anticipatory responses may reflect a trade-off between inhibiting and facilitating the onset of evidence accumulation via a gating mechanism as target appearance became more likely. Using a choice history analysis, we found that anticipatory responses were more likely following repeated choices, despite task randomization, suggesting that the balance between anticipatory and sensory responses was driven by a prediction of sensory timing. By fitting the gated accumulator model to the data, we found that variance in within-trial fluctuations in baseline evidence best explained the joint increase of anticipatory responses and faster sensory-guided responses with longer delays. Thus, we conclude that a prediction of sensory timing is involved in balancing the costs of anticipation with lowering the amount of accumulated evidence required to trigger saccadic choice, which could be explained by history-dependence in the baseline dynamics of a gated accumulator.

**New and Noteworthy:** Evidence accumulation models are used to study how recent history impacts the processes underlying how we make choices. Biophysical evidence suggests that the accumulation of evidence is gated, however classic accumulator models do not account for this, so little is known about how recent history may influence the gating process. In this work, computational modeling and experimental data from a free choice saccade task argue that predictions of the timing of sensory information are important in controlling how evidence accumulation is gated, and that signatures of these predictions can be detected even in randomized task environments where history effects may not be expected.

## Introduction

Pattern recognition is important for adaptive behaviour. Sometimes, the patterns we observe are representative of causal relationships in our environment, so we can use this knowledge to adapt the way we process sensory information and inform predictions about future events (Gershman et al., 2015; Griffiths and Tenenbaum, 2006; Richards and Frankland, 2017). Indeed, both empirical evidence and theoretical accounts point towards the importance of predictive feedback on even the earliest stages of sensory processing (Clark, 2016; Friston et al., 2006; Rao and Ballard, 1999). However, although predictions are often studied under conditions where they have an adaptive purpose, predictions may also be based on relationships that arise spuriously. In laboratory environments, it is common to randomize the presentation of sensory data such that information from one trial does not have predictive value for the subsequent trial. Despite this, people often show evidence for history effects, even when this confers no obvious advantage. While history effects in randomized tasks have been extensively studied in the psychological sciences (Abrahamyan et al., 2016; Bakan, 1960; Bar-Hillel and Wagenaar, 1991; Gilovich et al., 1985; Jarvik, 1951; LaPlace, 1951; Mookherjee and Sopher, 1994; Nickerson, 2002), less is known about how history effects influence the sensorimotor transformations that occur as we make decisions in real time.

Reaction times between the presentation of a sensory stimulus and the execution of a saccadic eye movement are thought to be determined by the time it takes to integrate sensory information from a baseline to a threshold (Bogacz et al., 2006; Busemeyer and Townsend, 1993; Carpenter and Williams, 1995; Dzhafarov, 1993; Ratcliff and Smith, 2004; Standage et al., 2014; Stone, 1960; Usher and McClelland, 2001; Wong and Wang, 2006). In decision-making tasks reported through eye movements, predicting the likelihood of one alternative favouring another has traditionally been accounted for through shifts in the baseline evidence (Bogacz et al., 2006; Carpenter and Williams, 1995; Glaze et al., 2015; Mulder et al., 2012; Ratcliff and McKoon, 2008), although a direct and unique link has been questioned (Fan et al., 2020; Simen, 2012; Urai et al., 2019). Reinforcement-learning approaches have since been applied to explain how predictions influence eye movements through trial-trial updating of baseline evidence (Bitzer et al., 2014; Kim et al., 2017), thus casting baseline evidence as a readout of a prediction that is updated on a trial-trial basis according to the outcome of a choice. However, baseline evidence is a fixed model parameter, aligned to the experimentally-controlled onset of a stimulus, that abstracts over the sensory processing that occurs prior to evidence accumulation (Schall, 2019) – it has no obvious relationship to a biophysical process. Thus, it is unclear how discrete trial-trial baseline updating in classic accumulator models is related to sensory processing that occurs prior to evidence accumulation, and how this may lead to the observation of discrete baseline evidence shifts in classic accumulator models.

Predicting the timing of future sensory information is informed by prior experience (Acerbi et al., 2012; Jazayeri and Shadlen, 2010; Taatgen and van Rijn, 2011). One natural timing problem faced in the evidence accumulation process is transmission delays between the onset of sensory information in the environment and its relay to regions thought to be involved in evidence accumulation, such as the frontal eye fields (Gold and Shadlen, 2001; Hanes and Schall, 1996) and the lateral intraparietal area (Huk et al., 2017; Shadlen and Kiani, 2013. In the visual system, it is generally known that different brain regions must adapt preparatory activity to account for variations in latency (Lee et al., 2016, 2020; Schmolesky et al., 1998; Shadmehr, 2010). Although timing the onset of the evidence accumulation process is an important component of decision-making (Teichert et al., 2016; van den Brink et al., 2021), any sensory processing that occurs prior to evidence accumulation is typically not addressed in models within the accumulator framework. An exception to this is the gated accumulator model of the visual-motor cascade in the frontal eye fields (Purcell et al., 2012; Schall et al., 2011), wherein transient responses to visual activity trigger the onset of the evidence accumulation. Thus, instead of describing the onset of evidence accumulation with a fixed baseline plus some sensory delay, the onset of the decision process is explicitly modeled as a consequence of upstream visual processing. However, this visual activity has been considered purely as consequent of feedforward visual activity, so the role of history effects are not well understood.

Here, we show behavioural data from a free choice saccade experiment that suggests a prediction of sensory timing is combined with delayed visual information to trigger the onset of evidence accumulation. To explain this, we developed a gated accumulator model, in which a continuously fluctuating baseline evidence signal is combined with delayed sensory information to trigger the onset of evidence accumulation. By explicitly modelling the delay interval with baseline fluctuations, the model captures expected reaction time distributions for multiple delay intervals concurrently, and can additionally account for anticipatory response distributions arising from baseline fluctuations in absence of external stimuli. To test how predictions derived from previous trials may be reflected in the gating process, we designed a free choice saccade task where participants had to balance withholding anticipatory responses with responding as fast as possible to one of two choice targets that were presented with variable asynchrony. A choice history analysis suggested that the outcome of the previous trial, despite randomization, influenced the balance between anticipatory and sensory-guided reaction times. By fitting the gated accumulator model to choice history data, we show that the gated accumulator could capture both anticipatory and sensory-guided reaction times, and that trial-trial updates to the variance in baseline dynamics best explained the influence of recent history on reaction times. This suggests that evidence accumulation during saccadic choice is gated by combining delayed sensory information with predictions from previous trials, even in randomized protocols, in order to adapt the timing of accumulation onset. Further, this suggests that variability in the baseline dynamics underlying gated accumulation may be subject to trial history effects.

## Methods

### Free choice experimental procedures

#### Participants

Six participants took part in the study. All had normal or corrected-to-normal vision. The participant gave written consent prior to undergoing the procedure. The study was conducted with an experimental protocol approved by Health Sciences Research Ethics Board, which adheres to the principles of the Canadian Tricouncil Policy Statement on Ethical Conduct for Research Involving Humans and the principles of the Declaration of Helsinki (2013).

#### Task

We adapted a free choice saccade task, first introduced by Schiller and Chou, 1998 and further studied in Noudoost and Moore, 2011; Soltani et al., 2013. The subject began by fixating on a central target projected onto a screen (20” Mitsubishi Diamond Pro [16x12inches] 1280x1024 pixels, 60Hz, contrast 75.2%, brightness 0%). Two targets, one in each visual field, were then presented with randomly assigned temporal asynchrony values (16ms, 33ms, 66ms, 99ms) and onset times drawn from *U* (750*ms*, 1250*ms*). The subject was instructed to look at either target, so long as it was done as fast as possible, without anticipating. The subjects were informed prior to the beginning of the experiment that the targets would come up at different times relative to each other. The subject was instructed to blink in between trials in order to minimize disruption of stimulus perception. The script for the task was written in MATLAB using Psychtoolbox.

#### Data acquisition and filtering

We collected a total of 80,550 trials from 52 sessions across our 6 participants. During each trial, eye movements were tracked via EyeLink 1000 Tower Mount (SR Research, Mississauga, Canada) with a 1000Hz infrared camera that tracked retinal position. Retinal position was calibrated prior to each session. The EyeLink apparatus was placed 60cm from the screen containing the saccade targets. Saccades were calculated o?ine using a saccade detection algorithm with a velocity criterion of 50°/s, and were individually verified through a custom visual marking program. Trials where the tracker lost the eye (either through tracking error or participant lapse) were excluded. Trials were also excluded if the eye position deviated more than 15 degrees from the horizontal. The first saccade following target onset, or the first saccade within 3 degrees of the future target position prior to target onset, were taken to calculate the saccadic reaction time and endpoint error. Saccades with a positive endpoint horizontal position were given a choice value of 1, and saccades with a negative endpoint horizontal position a choice value of 0.

#### Psychometric fitting

All data analysis was performed using custom scripts in Python. Psychometric functions were fit using the Psignifit toolbox with a Bayesian Inference fitting procedure (Kuss et al., 2005). We fit psychometric functions to a cumulative gaussian distribution. Choices occurring prior to 70ms (sensory delay cutoff) after target onset were discounted from psychometric fitting. To compare the slopes of the psychometric functions across choice sequences, we bootstrapped the marginal distribution that estimates the slope parameter, and used the position in the choice sequence as an ordinal variable to fit a linear regression model. The model therefore tests the hypothesis that there is a linear relationship between the slope of the psychometric function and the position in the choice sequence, against the null model that they are independent.

#### Statistical testing of differences in reaction times

Qualitatively, we observed an increase in specific portions of the distribution of reaction times as a function of choice history and delay intervals. From the perspective of our modelling framework, causal change in the underlying process may influence certain portions of the observed reaction time distribution without changing typical summary statistics such as the mean reaction time. Thus, analysis of variance tests were not appropriate to analyze effects that may not be manifest as changes to mean reaction times. In order to avoid this issue, we chose to take the standard deviation as a metric that would be sensitive to changes to anticipatory and express saccades. To estimate of the confidence interval around the standard deviation, we bootstrapped each reaction time distribution 1000 times and computed the 95% confidence interval of the standard deviation. We then compared each successive choice sequence and determined if the intervals overlapped.

#### Fitting to free choice saccade data

We fit a gated accumulator model (Results: Gated Accumulator) to reaction time data from the free choice saccade task. We used differential evolution, a stochastic optimization technique in the *Scipy* optimization toolbox, to fit a parameter set for each delay bin. This algorithm allowed us to avoid local minima in the parameter space that interfere with gradient-based methods. We used the ‘best1bin’ search method, with a maximum of 1000 iterations performed, population size of 15, recombination constant of .7, and convergence tolerance of .01. The 5 parameters of the gated accumulation model were given the following boundary conditions

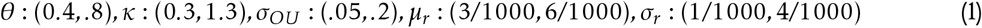

Before fitting, 5 alignment points for delay times were chosen (evenly spaced between 750 and 1250ms), and reaction time data was discretized into these 5 bins relative to the delay time (first target onset). Thus, each dataset contained 5 overlapping reaction time probability distributions. A model was then generated with a target onset time at each of these 5 delay times. To obtain a model error, the Kullback-Leibler divergence between each delay bin (model vs data) was summed across the 5 delay bins. It was this summed error that was used as a cost function for the differential evolution algorithm.

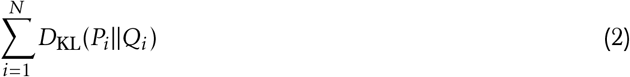

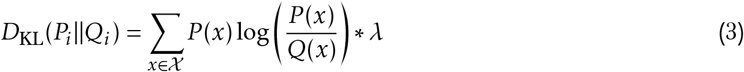

where *i* is the delay bin start, generated by *N* equal spacings between 750 and 1250 ms, *P*_*i*_ (*x*) is the reaction time model probability for delay bin *i* and reaction time bin *x, Q*_*i*_ (*x*) is the same for the data probability distribution, and *λ* was a constant scaling term to prevent numerical underflow. The discrete model probability was calculated from the full probability density function by sampling 10,0000 times from the continuous model over the same set of 50 discrete reaction time bins as the data histogram was computed from.

## Code and data availability

Python code for the gated accumulator model, data analysis, and model fitting can be found on https://github.com/BJCaie/Gated-Accumulation. Data is available at https://osf.io/6zjkd/

## Results

We designed a free choice saccade task (*Methods: Free choice experimental procedures*) where participants had to rapidly respond to the onset of sensory information while withholding anticipatory responses. Each trial begins with a central fixation period drawn from a uniform distribution between 750 and 1250 ms (Fig 1A). This period was followed by the appearance of two visual targets, one in each visual hemifield. The targets appeared at different times relative to each other, with the target onset asynchrony (TOA) randomized across 9 discrete values 0, +/- 16, 33, 66, 99ms. Positive asynchrony values will denote the right target appearing prior to the left, and negative asynchrony values will denote the left target appearing before the right. Participants were instructed to direct a saccade to either target as fast as possible, without anticipating. Although both delay and TOA were randomized, any given sample of trials from a uniform distribution may have biased statistics in the spatial/across-trial (target location) and temporal/within-trial (target onset and asynchrony) domain (Fig 1C-D).

**Figure 1.**
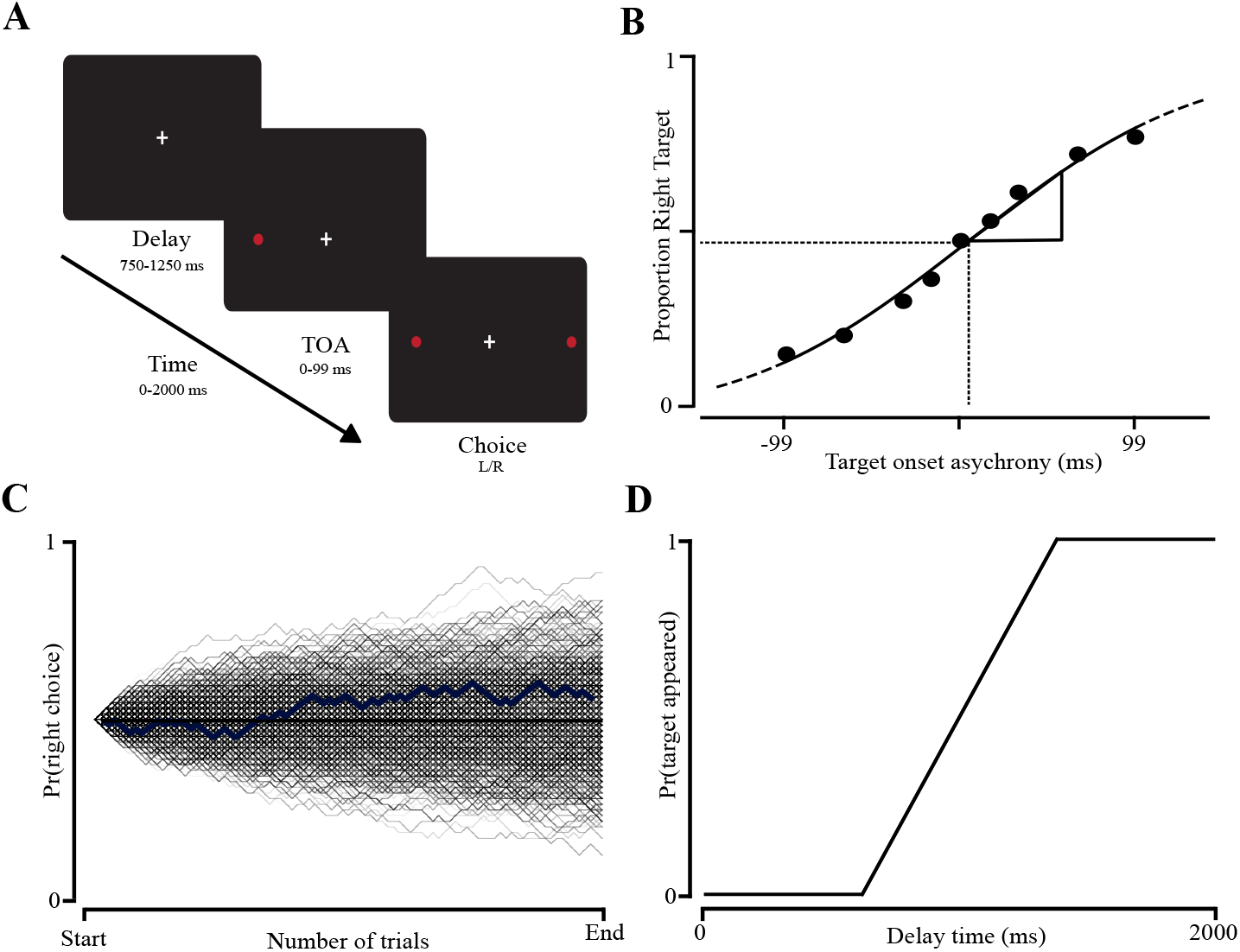
Free Choice Saccade Task. **A**: Participants directed saccades to either of two free choice targets as fast as possible, with a randomized delay time and target onset asynchrony (TOA).**B**: Group-average psychometric function, depicting the probability of right target (y-axis) as a function of TOA. **C**: Across-trial task statistics. Depicted is a random walk (left), showing the probability of right target selection, beginning at 0.5, across different hypothetical blocks. **D**: Within-trial task statistics. Depicted is the probability of the first target appearing as a function of the delay time of the trial.

### Choice probability is history-dependent

Psychometric analysis revealed that choices are probabilistic with respect to the magnitude of asynchrony (Fig 1B). Choice behaviour binned by asynchrony indicated a progressive increase in rightward choices when the right target appeared earlier. The binned choice data was fit to a sigmoidal curve with a psychometric fitting procedure using Psignifit (Schütt et al., 2016). Note in this analysis only choices occurring after the target onset plus a minimum delay of 70ms were included, as is often standard when excluding anticipatory trials (Nijhawan, 2008; Schmolesky et al., 1998). Synchronous targets resulted in choices closest to chance with respect to left or right direction, while a target appearing well before another increased the likelihood of response towards it. This indicated that participants were aware of the task instructions, and that the range of TOAs used in the task were sufficient to evoke a change in choice direction probability.

In randomized tasks, trials are uncorrelated, so analysis of behaviour often rely on treating trials as statistically independent. However, it is common to infer patterns in short sequences of randomized information (Jarvik, 1951; Nickerson, 2002). In our task, there is no inherent advantage to relying on information from past trials, but each choice trial is embedded in a unique sequence. A binary choice tree for 5 trials in the free choice task is schematized in Fig 2A. For future analyses, we compared sequences of choice repetitions to the same target direction, and sequences of choices alternating targets, as measures of choice sequences with a stereotyped local structure that may induce bias away from the true generative statistics of the task (Cho and Cohen, 2002). Fig 2B-C shows a histogram of the number of repetitions and alternations in the present data-set. Because the data become increasingly limited with higher sequence number, analyses were conducted up to a sequence length of 5.

**Figure 2.**
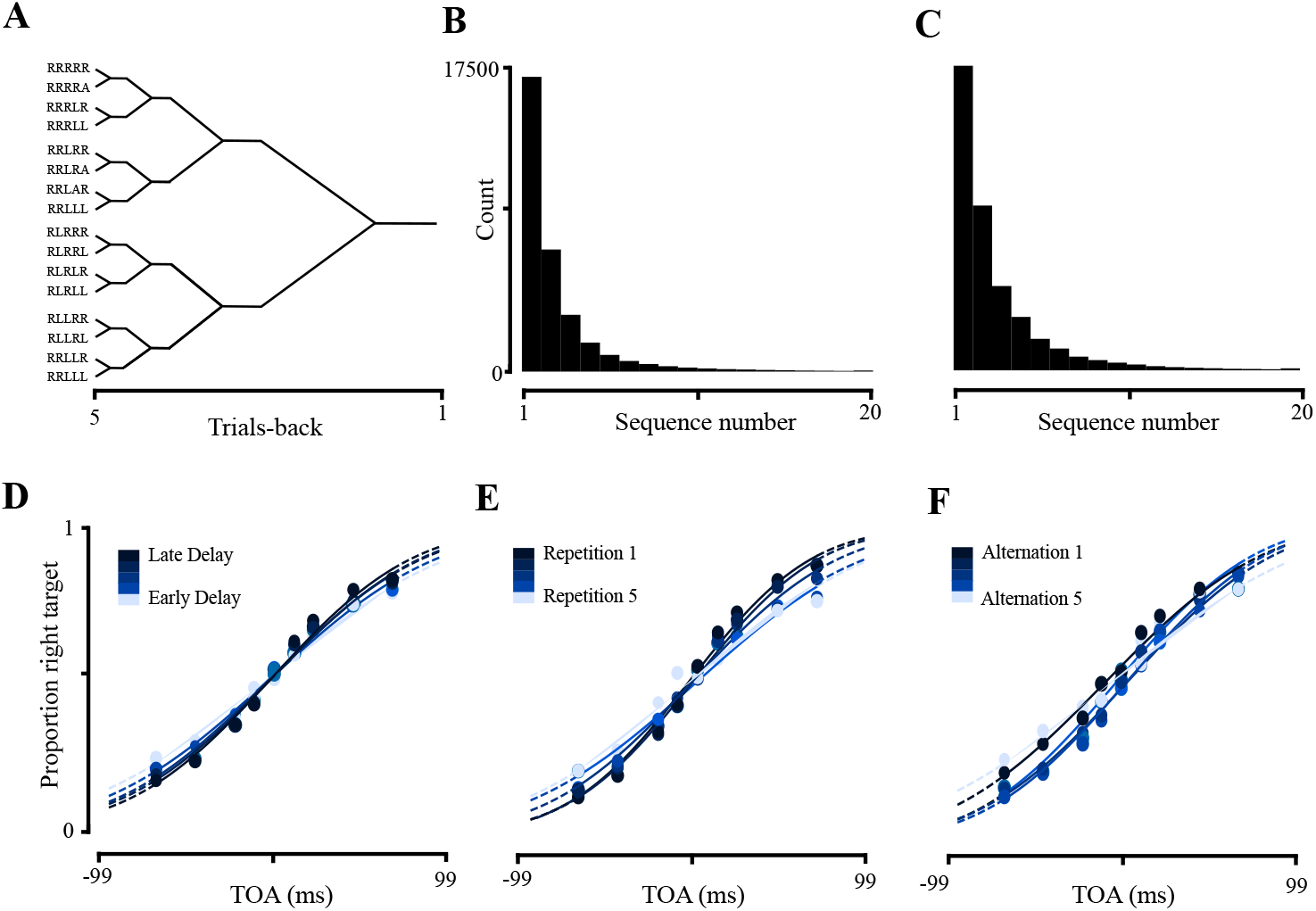
Choice history and delay time influence choice probability. **A**: Choice history tree depicting the possible alternatives of a binary choice up to 4 trials back. **B**: Histogram of the number of choice repetitions across all participants. For each sequence number, a single tally was recorded if the same choice direction had been chosen for that many trials previously. **C**: The same as B, but for the number of successive choice direction alternations. **D**: Group-average psychometric functions with respect to TOA, where the data has been split into 4 delay bins equally spaced between 750 and 1250 ms. The colour gradient denotes the delay period, low to high. **E**: Psychometric functions split into the number of previous choice repetitions. Here, each repetition sequence was given an independent repetition score, such that no trials in one repetition set are contained in another **F**: The same as **E**, for choice alternations.

Psychometric analysis (*Methods: Psychometric Fitting* ) revealed that choice probability was influenced by the number of previous repetitions and alternations. In Fig 2E, psychometric fits to group-averaged repetition data are plotted as a function of repetition number (blue gradient), showing a progressive decrease in the slope of the sigmoid function. We interpret this as indicating that following choice repetitions, participants relied less on the TOA of the current trial, and more on information derived from past trials. To quantify this, we estimated parameters of the psychometric function using a Bayesian procedure (*Methods: Psychometric fitting* ). We then bootstrapped estimates of the slope from the resulting probability distribution over the slopes, and performed a Kruskal-Wallis to ask if the parameter estimates came from the same distribution (*F* = 2829.13, *p* = 0). This suggests that the distribution of best-fitting psychometric slopes changed across repetitions. We performed this same analysis for sequential alternations, finding that choice probability was also not constant (*F* = 476.23, *p* = 9.18^−102^).

In addition to the TOA, we also assessed the role of the delay interval. We next assessed whether the length of the delay interval influenced choice probability. To do this, we performed the same binning procedure as previously outlined for choice sequence, but instead we binned the delay period into 5 equally spaced intervals between 750*ms* and 1250*ms*. The resulting psychometric functions are shown in Fig 2D. Again, we performed a Kruskal-Wallis test, which supported the qualitative observation that the bestfitting psychometric functions were not constant when separating data by the length of the delay interval (*F* = 384.77, *p* = 5.43^−82^). In combination, these analyses support the conclusion that both choice sequence and delay time influenced choice behaviour.

### Choice history jointly influences anticipation and the accumulation of choice evidence

Analysis of reaction time distributions provides a window into the processes underlying saccade triggering (Carpenter and Williams, 1995; Hanes and Schall, 1996; Schall, 2019). In order to better understand how the choice process evolved throughout the trial, we therefore performed a reaction time analysis of the free choice saccade task based on the delay interval and previous trials. Saccades are often grouped into three classes based on reaction time: anticipatory responses that occur before stimulus onset, sensory guided responses, and an ‘express saccade’ regime that lies between the two (Kingstone and Klein, 1993). In the psychometric analyses, only choices occurring after sensory onset plus a minimum delay of 70ms were considered (Schmolesky et al., 1998), so as to analyze the role of target onset asynchrony. However this was only a subset of the total behaviour; while participants were asked to try not to anticipate, they were also asked to go as fast as possible, so in a subset of trials participants responded prior to target appearance.

First, we assessed the influence of the length of the delay interval on anticipatory and sensory-guided responses. Fig 3 plots the cumulative distributions of reaction times split into 5 equally sized delay bins between 750 and 1250ms. We aligned the distributions in two ways. First, we aligned relative to the onset of the target, and plotted anticipatory responses as negative time relative to this (Fig 3A). Next, we aligned the data to the onset of the trial, such that each delay bin will have a mixture of onset times bounded within the delay bin (Fig 3B). In each view, it can be seen that anticipatory responses increase, and sensory-guided responses get faster, when the delay interval was longer (Reddi and Carpenter, 2000; Standage et al., 2014). Qualitatively, this resulted in a broadening of the distribution as a function of the delay interval, since an increase in anticipatory responses corresponds with an increase in the early tail of the distribution. To quantify this, we randomly sampled from each distribution and bootstrapped the empirical standard deviation, and computed the 95% confidence interval for each. This supported the conclusion that reaction time variance was dependent on the delay interval.

**Figure 3.**
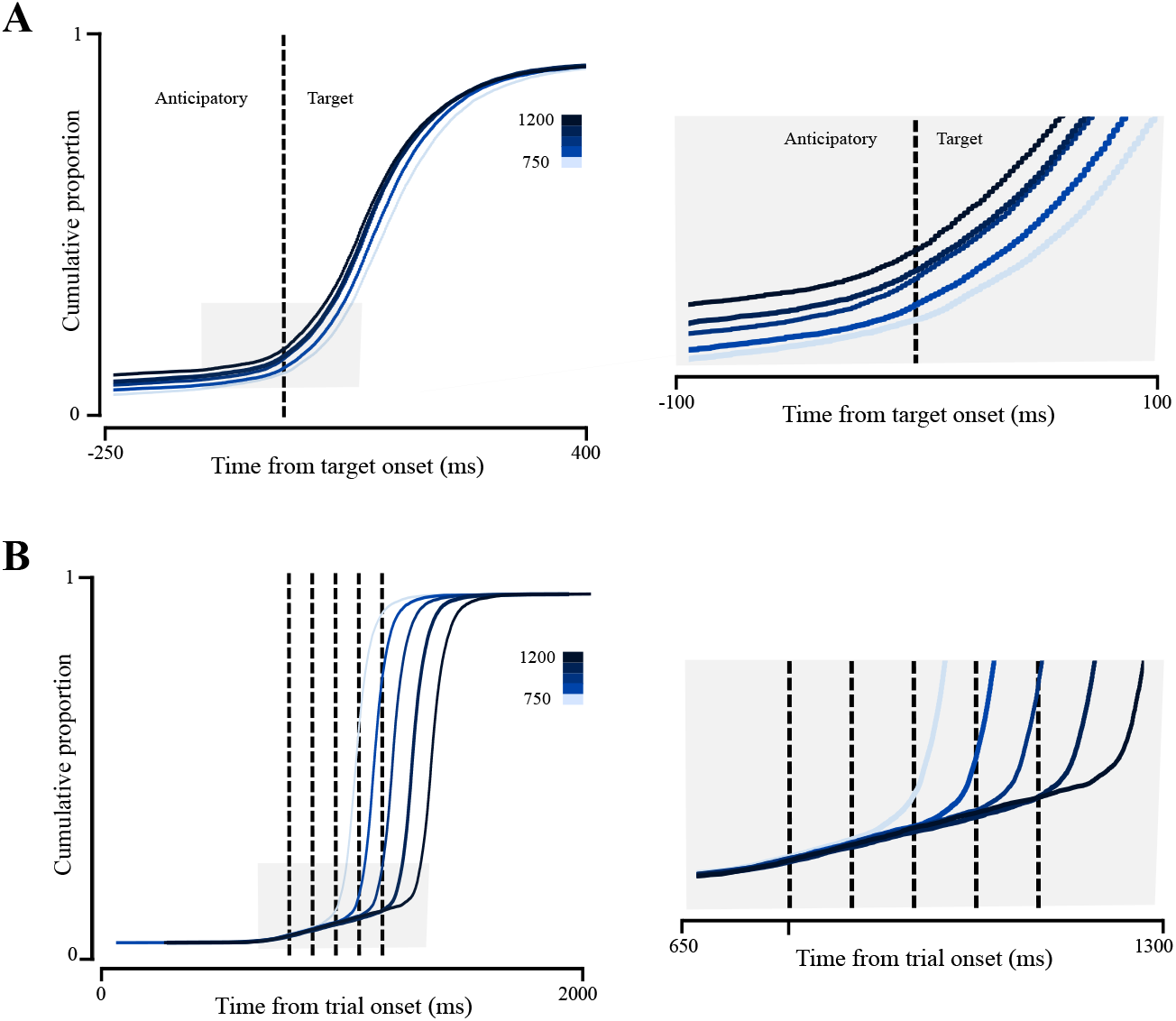
Anticipatory and sensory-guided reaction times are influenced by delay time. **A**: Cumulative probability of reaction times as a function of 10 delay bins, aligned to the onset of the target (left) and trial (right). The onset of the cumulative probability functions deviating from zero in target-aligned responses represent the proportion of anticipatory responses visible in the trial-aligned graph. **B**: Cumulative response functions separated by the number of previous repetitions of the same choice direction. **C**: The same, for alternations.

We next analyzed reaction times as a function of choice history (sequential repetitions and alternations). Fig 4 plots the reaction times aligned to the target dependent on how many previous repetitions (Fig 4A) and alternations (Fig 4B) preceded the current trial. Here, it can be seen that sequential repetitions were associated with an increase in anticipatory responses, as seen in the cumulative proportion lying before the dashed line indicating target onset, and faster post-target responses. However, this speed advantage was negated for responses in the median range, and even reversed for later responses, consistent with the idea that choice sequence influenced an association with baseline evidence in accumulator models (Nakahara et al., 2006; Noorani, 2014). This effect was detectable but noticeably diminished in magnitude for alternations, consistent with the idea that sequence effects may operate over different timescales when targets alternate (Cho and Cohen, 2002). In summary, our results suggest that reaction times were jointly influenced by choice history and the delay time of the trial. By analyzing reaction times not by summary statistics such as the mean or median, and instead viewing entire distributions relative to different onset times, we were able to conclude a joint influence of delay time and the previous trial on early and express saccade responses.

**Figure 4.**
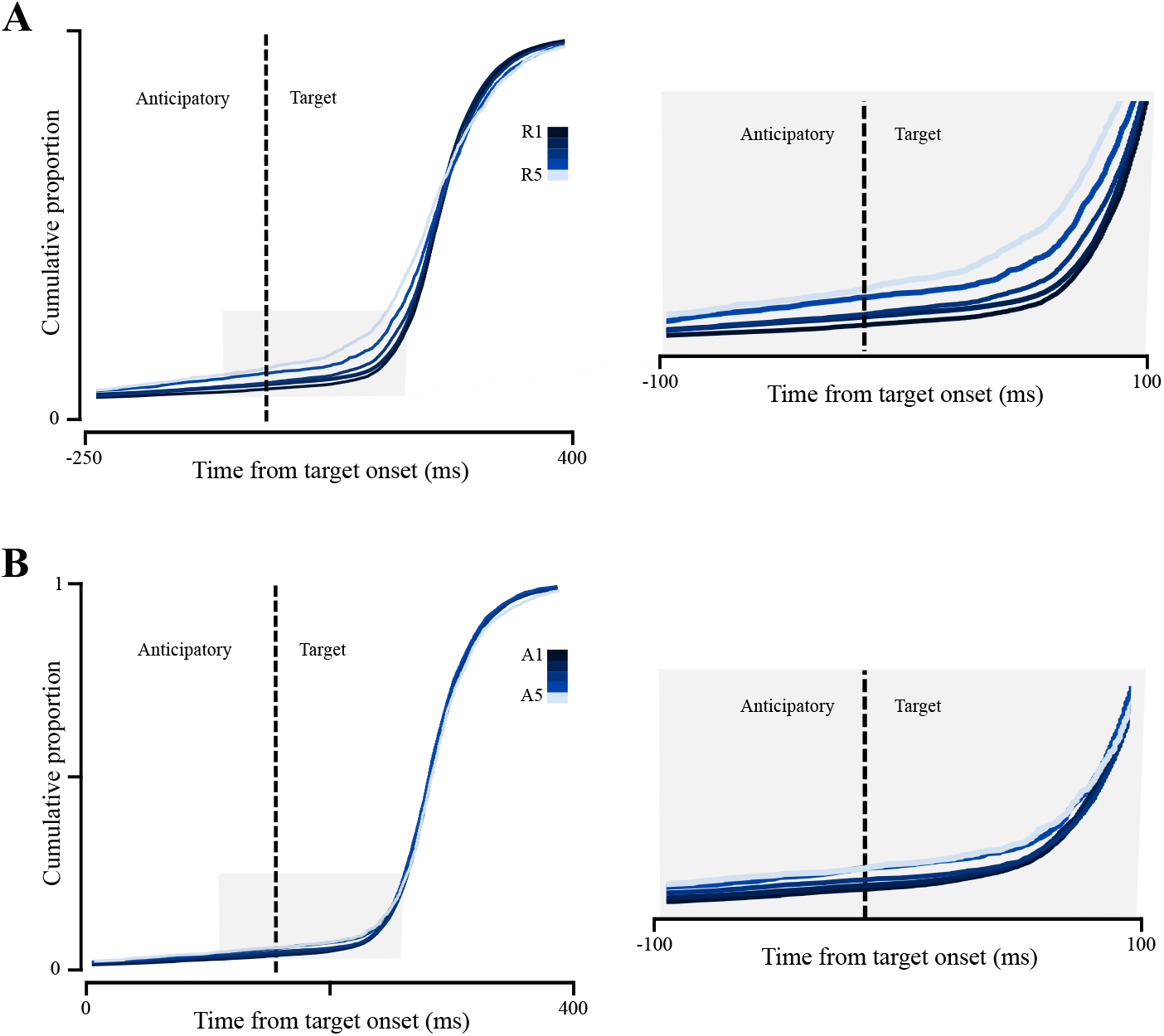
Anticipatory and sensory-guided reaction times are influenced by choice history. **A**: Cumulative probability of reaction times as a function of 10 delay bins, aligned to the onset of the target (left) and trial (right). The onset of the cumulative probability functions deviating from zero in target-aligned responses represent the proportion of anticipatory responses visible in the trial-aligned graph. **B**: Cumulative response functions separated by the number of previous repetitions of the same choice direction. **C**: The same, for alternations.

### Modelling saccade triggering as a gated accumulation

The gating of inputs to different neural structures is an established mechanism of information transfer in the brain (Gisiger and Boukadoum, 2011). In Purcell et al., 2012, the accumulation of evidence favouring one of several choice alternatives is hypothesized to be gated below some threshold for saliency, allowing for the selective accumulation of information in the presence of a continuous stream of changing visual scenes. It is commonly thought that such evidence is integrated or amplified towards a threshold, upon which a corresponding motor command is produced (Cisek et al., 2009; Schall, 2019). Models of the accumulation process have served a dual role, bridging an algorithmic explanation of the heavy-tailed distributions characteristic of reaction times with a mechanistic explanation of the delayed ramping activity of single neurons in regions of the brain such as the frontal eye fields (Gold and Shadlen, 2007; Hanes and Schall, 1996). In recognizing both the necessity and empirical evidence for the control of the accumulation process by a gating mechanism, Purcell et al., 2012 extended the descriptive capacity of accumulator models to explicitly include the visual transformations occurring prior to the onset of evidence accumulation, thereby replacing the discrete baseline evidence of traditional accumulator models (such as the drift-diffusion model, LATER model, and linear ballistic accumulator) with a continuous visual transformation.

In this section, we derive a gated accumulator that receives a continuous input to an evidence accumulator during the delay interval of a trial. We first decompose the model into two sections: a section explaining the *baseline dynamics*, or how baseline evidence fluctuates throughout the trial prior to the onset of sensory information, and a section explaining the *evidence accumulation*. Then, we explain how these processes may be linked by a gating mechanism, and how we combine the two processes algorithmically to obtain a single probability density for reaction times.

### Baseline dynamics

We first describe the baseline dynamics (Fig 5) that evolve throughout the delay interval of a trial prior to the onset of a visual stimulus. We chose an Ornstein-Uhlenbeck (OU) process to model the baseline dynamics for the free choice saccade data. This was chosen because it is a well-characterized dynamical system with tractable properties that can describe the evolution of a stochastic process from a baseline value to a steady-state. However, it should be noted that this does not constitute an argument for the necessity of an OU-like process in the neural processes underlying gated accumulation, merely its sufficiency to describe the data. The OU process is a stochastic differential equation that describes the time evolution of a random variable changing proportionally to the distance from an expected mean (*Fig 2*). Let *B*_1_, *B*_2_, …*B*_*n*_ be a set of processes which evolve as a time series during a trial of length *T*_0:*T max*=2000_, where *b*_*i*_ = *b*_1_, *b*_2_, …*b*_*n*_ is a single process satisfying the stochastic differential equation

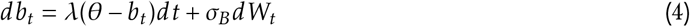

where *λ* is the drift rate, *θ* is the steady-state baseline the process reverts to as time approaches infinity, and *W* is a Wiener process scaled by *σ*_*B*_.

**Figure 5.**
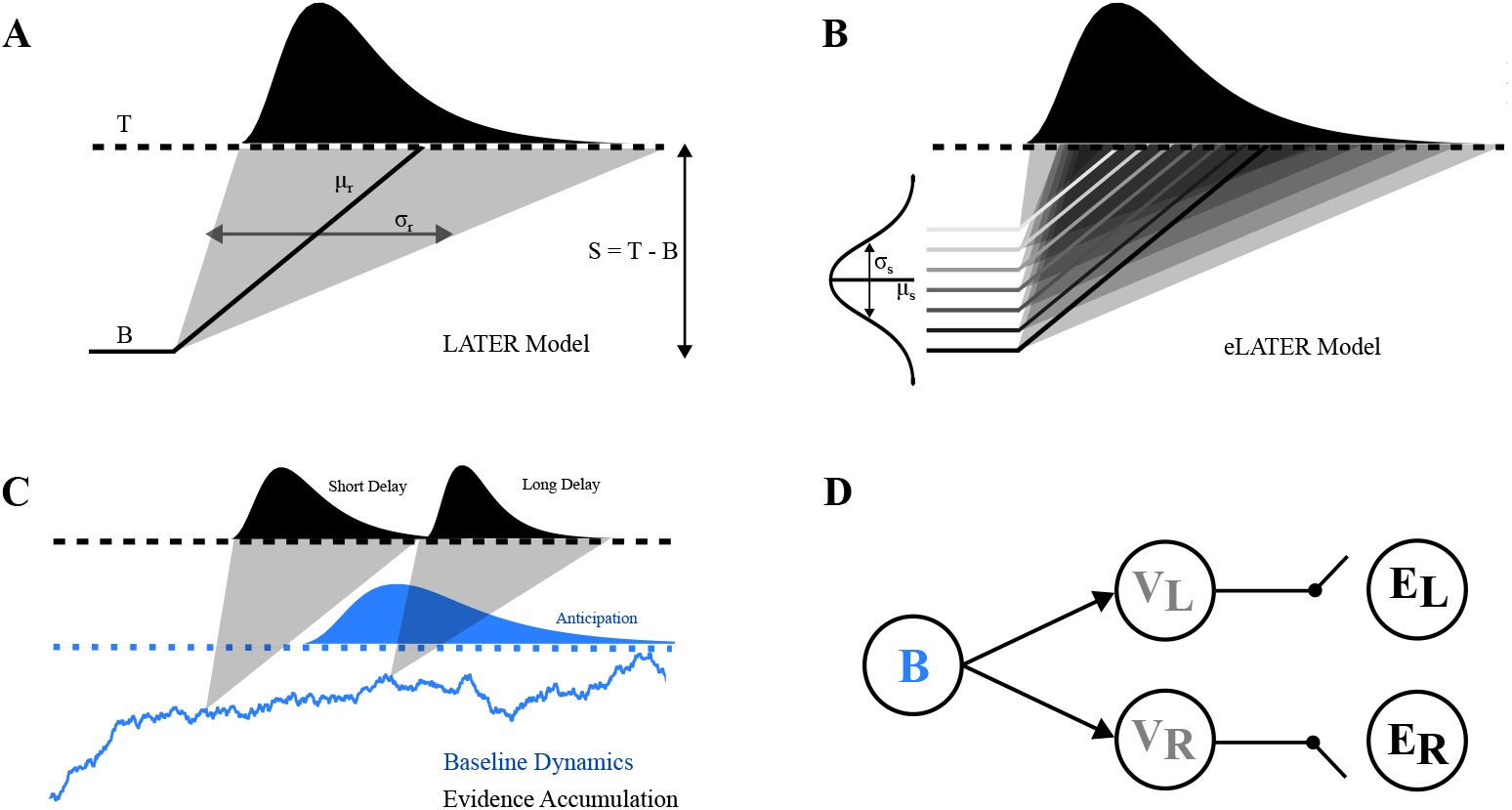
Gated accumulation of choice evidence. **A:** Schematic of a linear approach to threshold with ergodic rate (LATER) model. Sensory evidence is integrated upon a baseline level of choice evidence towards a movement-triggering threshold. The strength of sensory evidence typically controls the average rate of rise. Variability in the rate of rise yields variability in reaction times (top curve). Time in the model begins relative to the onset of a repeated stimulus **B:** Extended LATER model (eLATER), with variance in the baseline giving rise to a distributed integration process. The resulting probability density function is integrated over the threshold-baseline distribution. **C:** Gated accumulation. Baseline dynamics, governed by an Ornstein-Uhlenbeck process, are combined with the delay time of a given trial to yield a joint distribution of anticipatory and sensory-guided reaction times. **D:** Circuit diagram of a two-alternative gated accumulation process, with the baseline dynamic *B* combined with visual transients *V* to trigger evidence accumulation *E*.

We further assume that if this system reaches the expected baseline *θ* prior to the onset of a choice target, then an anticipatory responses is triggered. Thus the rate *λ* dictates if and how close to *θ* the system reaches prior to target onset, and thus if an anticipatory response is triggered. The anticipatory response distribution is thus defined by the first-passage problem where the threshold is equal to the long-run mean *θ*, which is described in Lipton and Kaushansky, 2018 as the solution to the problem

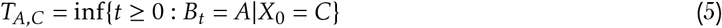

with change of variables

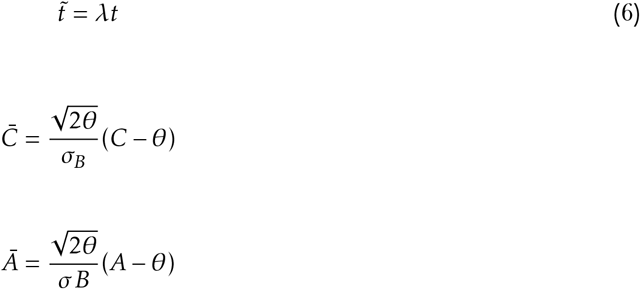

The anticipatory density function for the case where *Ā* = 0 is known.

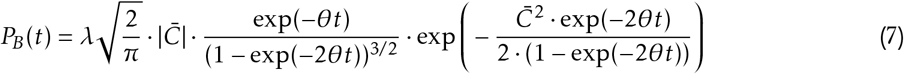

Because *Ā* = 0 is equivalent to setting the threshold crossing *A* to equal *θ*, this equation becomes the first passage time solution for when the threshold *A* = *θ*. Thus, this equation describes the probability density of how long it takes the OU process to reach the long-run mean *θ*, scaled by the rate parameter *λ*.

A graphical depiction of the baseline dynamics of a sample OU process is depicted in Fig6). In panel *A*, different instances of the same OU parameter set are plotted in the blue traces, with each evolving according to the difference between its current state and a mean reversion parameter *θ* = 1.5 at a rate according to *λ* with variance *σ*_*B*_. We can solve analytically for the first passage time for a given parameter set, yielding our anticipatory response function, as the time it takes to reach the steady state (*Eq. 4*). Additionally, we have a Gaussian probability density of baseline activity at each time point throughout the delay period (Fig6C), which we will use to model a distributed evidence integration process with variability in the baseline, as in Nakahara et al., 2006.

**Figure 6.**
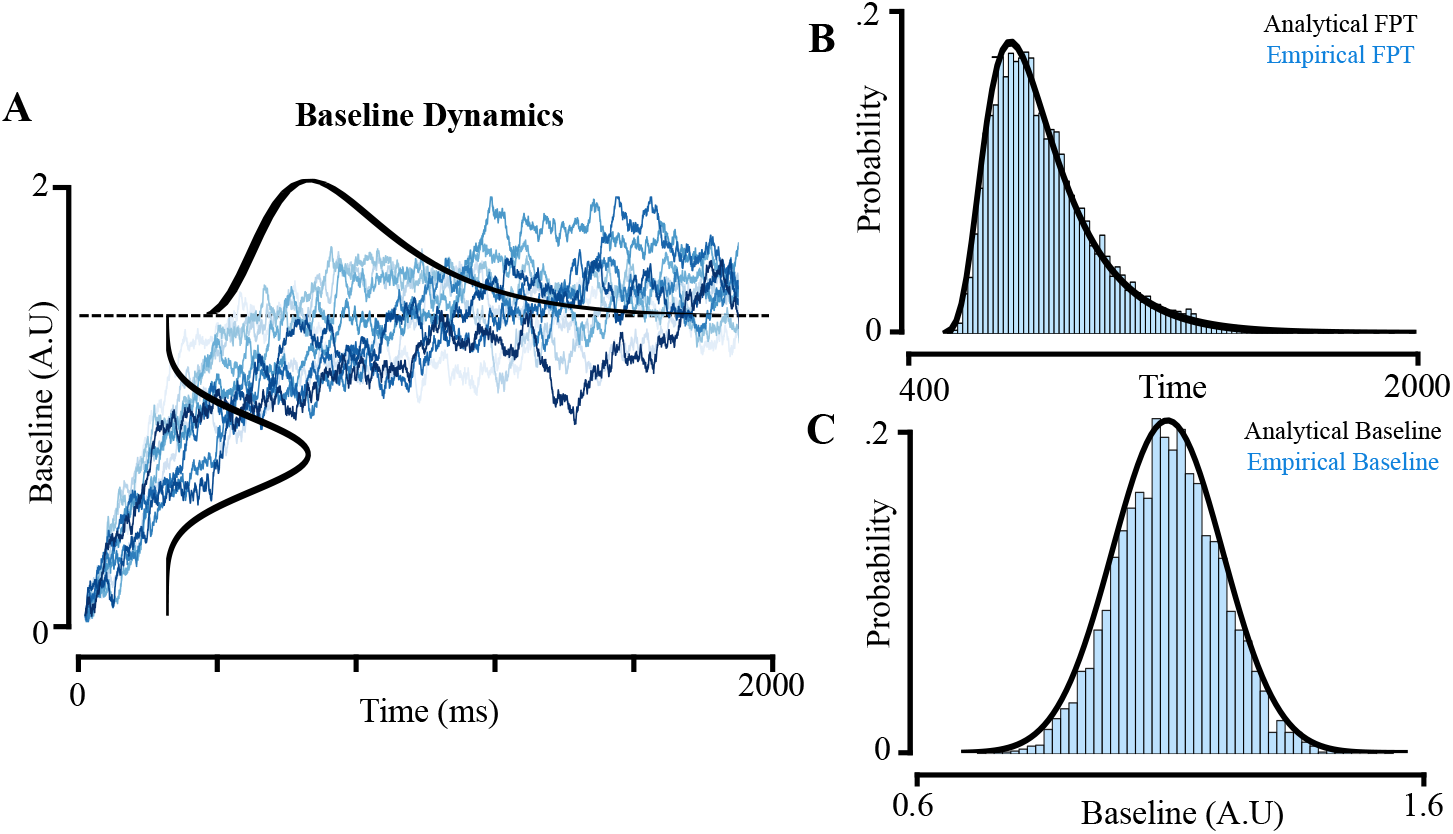
Baseline Dynamics. **A**: Sample paths of an Ornstein-Uhlenbeck process used to model the baseline dynamics. Each blue trace is one sample path drawn from the same parameter set: *θ* = 1, *µ* = 1.5, *σB* = 0.5, X0 = 0 **B**: The first passage time for this parameter (black: analytical solution, blue: empirical histogram for 1000 processes) set is plotted as the probability that the OU process has reached the *µ* value at time t. This is also seen in the heavy-tailed distribution above the threshold in **A. C**: probability distribution of baseline values at t = 250 ms. The distribution at any time follows a Gaussian distribution (colour scheme same as **B**).

### Evidence accumulation

We next describe the integration of evidence to a threshold following the onset of sensory information. For simplicity, we chose to describe the evidence accumulation process that occurs after the onset of a choice target as minimally as possible, so as to focus on the effects of input during the delay period. A simple and interpretable evidence accumulation model is the linear approach to threshold with ergodic rate (LATER) model (Carpenter and Williams, 1995; Carpenter, 1981; Noorani, 2014). The LATER model captures reaction time as a deterministic signal with variability across trials using 3 free parameters. Any single trial is determined by

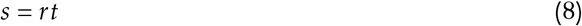

where *t* is the response time, *s* is the threshold-baseline difference, and *r* is the rate of rise. The distribution of reaction times can be derived using process parameters *µ*_*r*_ , the mean rate of rise, *s* the distance from baseline to threshold, and *σ*_*r*_ , the variance of the rate of rise

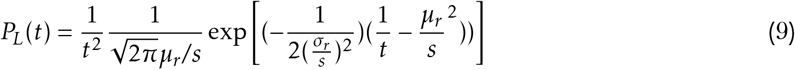

In the LATER model, the baseline is a single value, and the variance required to explain a *distribution* of reaction times comes from trial-trial variance in the integration rate. Although the LATER model fits the majority of a reaction time distribution, it does not account for anticipatory responses and the short latency reaction times typical of eye movements, often termed ‘express saccades’ (Noorani, 2014). In order to treat the entire reaction time distribution under this framework, and crucially the responses close to the time of target onset, a derivation of the LATER model with variance to the baseline value was proposed (Nakahara et al., 2006). This ‘extended LATER’ model has an analytical solution for the reaction time probability density if the prior is Gaussian, *N* (*µ*_*s*_*σ*_*s*_)

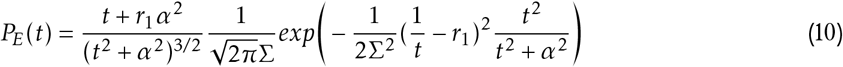

where 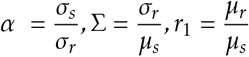.

The extended LATER model allows us to calculate reaction time distributions for different *distributions* of baseline activity, rather than using the single baseline parameter in the LATER model. Because of this, we can then model the expected reaction time distribution for different delay lengths by combining the extended LATER model with a time-varying baseline dynamic.

### Combining evidence accumulation with baseline dynamics via gating

We next describe how the two previously described probability density functions, the anticipatory response distribution *P*_*B*_ derived from the baseline dynamics, and the evidence accumulation distribution *P*_*E*_, are combined into a common process with a single probability density function. We first treat the simplest case, where only a single target onset time is used. Thus, we need to compute the anticipatory first passage problem for the *OU* process, as well as the mean and variance of the process itself at the corresponding delay time, which we will use as the mean and variance to the baseline Gaussian distribution in the extended LATER model. We can thus compute the mean of the extended LATER baseline *µ*_*r*_ at time *t* as the expectation of the Ornstein-Uhlenbeck process, which is well-known

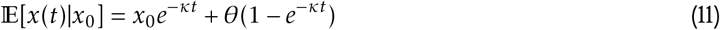

and the covariance

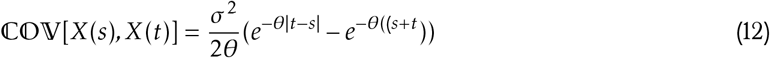

where *s* = 0 and *t* = *y*. In order to convert the rising baseline of the OU process into a threshold-baseline distance for the extended LATER model (where a smaller value indicates a shorter distance and thus a faster time to threshold and reaction time), we translate *µ*_*r*_ as

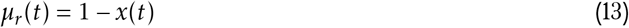

where *µ*_*r*_ is bounded between 0 and 1. Thus, while adding 4 free parameters from the OU process, we reduce two free parameters from the extended LATER process *pT* by allowing the OU process to control the baseline Gaussian at a given target onset time *t*. We can then define the gating process, *G*, as a combination of the anticipatory first passage distribution *P*_*B*_, and the extended LATER solutions *P*_*L*_ for a distribution of delay intervals *D* over the interval [*a, b*].

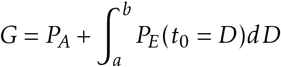

A key distinction between the way in which we have derived this model and other evidence accumulator models is that by explicitly accounting for the delay interval and anticipatory responses, the onset of sensory information is no longer a fixed, experimentally controlled parameter that references *t = 0*, or *t = 0* plus some fixed sensory delay parameter. Instead, the model output is the combination of the parameters of the model and the chosen delay times.

Fig7) depicts a cartoon scenario illustrating the impact of the delay time of a given trial on the outcome of different baseline dynamics. If two different baseline dynamics, depicted in dark and light blue respectfully, were aligned relative to the onset of a stimulus but had different delay times (Fig7A), they may produce identical baselines at the time of target onset, and as such not be dissociable. However, if the same baseline dynamics were shifted such that the common starting time was the trial, and not the target, the same baseline dynamics would lead to an observable difference in reaction times (Fig7B). This highlights that distinctions between underlying processes can be masked by the way in which reaction time data is aligned, while identical processes can result in quantitatively different behaviour if the temporal statistics of the task are appropriately varied.

**Figure 7.**
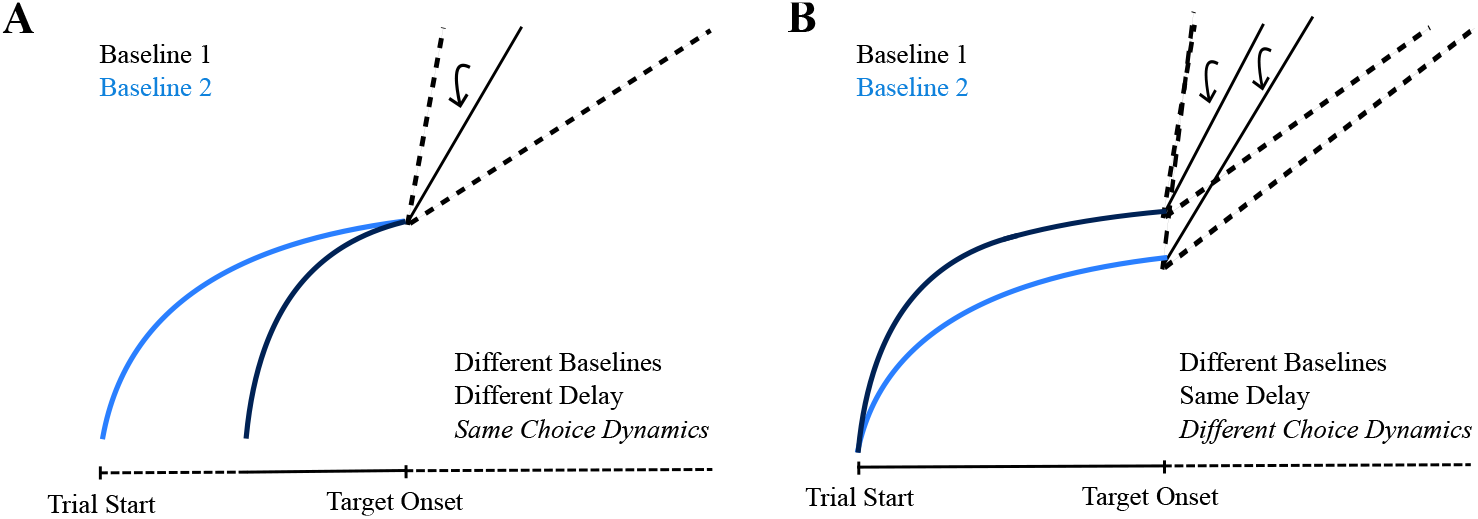
Inferring difference in baseline dynamics depends on starting point. **A**: two trials where baseline dynamics (high rate parameter: light blue, low rate parameter: dark blue) with the same delay time reach the different baseline levels, resulting in distinct accumulation processes. **B**: two baseline dynamics with different delay interval lengths, resulting in identical baseline evidence at target onset, and thus indistinguishable reaction time distributions.

### Gated accumulation of sensory timing expectations accounts for choice history and anticipation

Biophysically motivated models of evidence accumulation have suggested that the onset of accumulation is gated in order to prevent prepotent responses to the continuous stream of sensory inputs we are subject to in natural visual scenes (Purcell et al., 2012; Schall et al., 2011). We derived a model in which the gating of evidence accumulation is not driven solely by delayed sensory information, but also by within-trial fluctuations in baseline evidence dynamics. Here, we show that trial-trial changes in the baseline dynamics can account for the dependence of reaction times on choice history that was observed empirically.

To do this, we fit the group-averaged reaction time data to the gated accumulator model, split up by choice repetitions so as to isolate the strongest observed effect of choice history (Methods: Fitting to Free Choice Saccade Data). The model fitting procedure yielded a joint probability density function for reaction time data continuously throughout the trial period, which was post-hoc binned into 5 delay quantiles for visualization. Fig8) depicts the result of the model fitting. In Fig8A), the baseline dynamics for the best fitting model are shown (the mean baseline is shown in the black trace, with the shaded blue depicting the standard deviation of the baseline at each time point, and the dotted line depicting the threshold).

**Figure 8.**
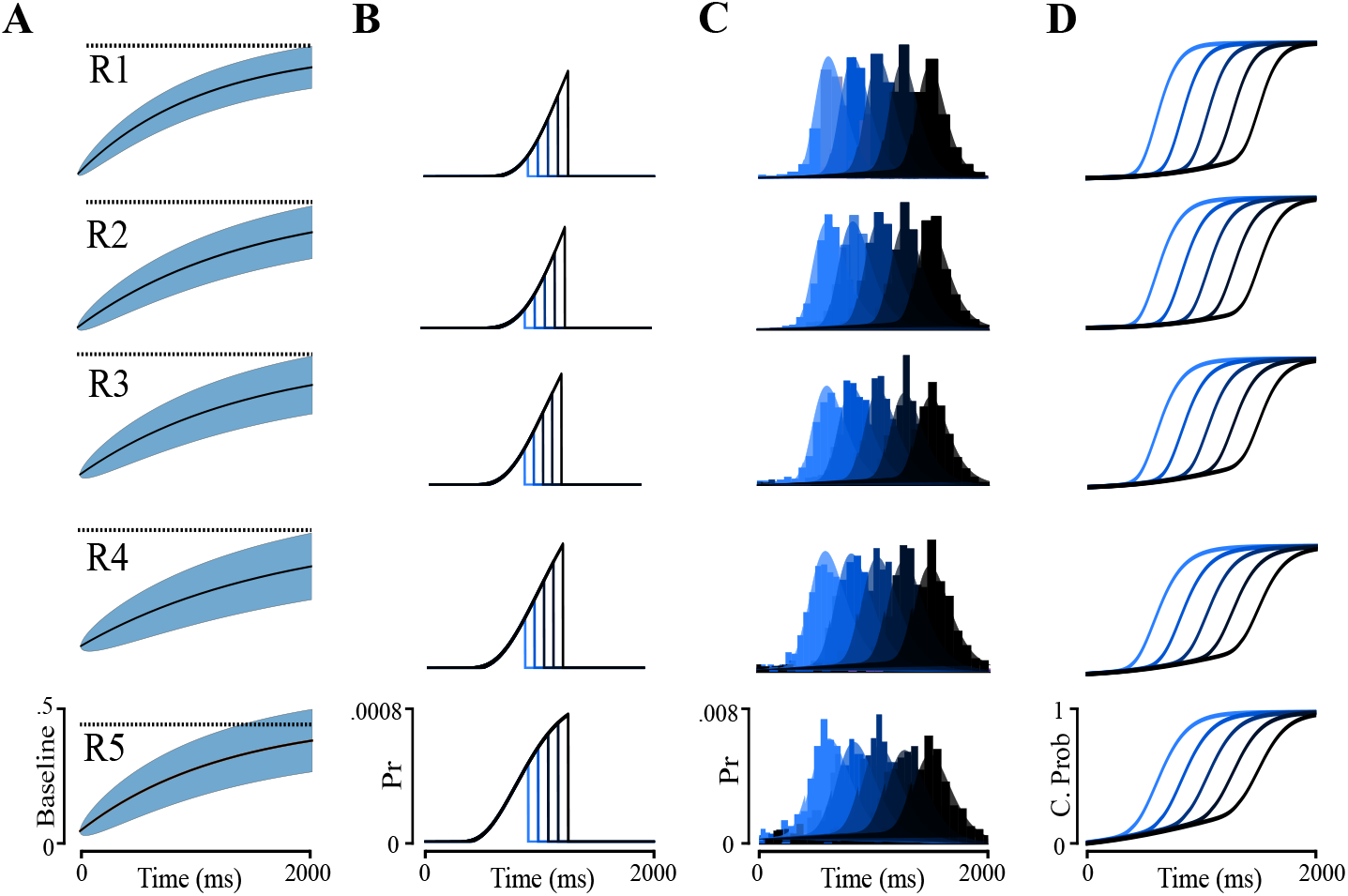
Gated accumulation explains dependence of reaction times on choice sequence and delay time. **A:** Best fitting baseline dynamic for each dataset, where each row denotes the fit to repetition 1 (top row) through 5 (bottom row). The mean of the baseline dynamic is plotted in black, and the variance is plotted in the shaded blue. The dotted line indicates the value of the mean-reversion parameter *θ*. **B:** Anticipatory first passage of the model for 5 delay times, each cut off at target onset. **C:** Model reaction time density functions (shaded curves) are plotted against the data (histograms). **D:** Cumulative density functions from the best-fitting model.

Here, a dependence of the model fitting results on the choice repetition sequence on the variance of the baseline dynamic *σ*_*B*_ was observed. There was a progressive increase in *σ*_*B*_ as repetitions increased (*R*_1_*σ*_*B*_ : .112, *R*_2_*σ*_*B*_ : .119, *R*_3_*σ*_*B*_ : .127, *R*_4_*σ*_*B*_ : .157, *R*_5_*σ*_*B*_ : .183,), while no consistent trend was observed in the other baseline dynamics parameters *θ* or *κ*, nor the sensory integration parameters *µ*_*r*_ or *σ*_*r*_ . This indicates that the trial-trial changes observed in the reaction time data may not be best explained by shifts in the baseline *per se*, but the variance in the process that gives rise to its average value. As a result, the model best explains the data by a combination of increased anticipatory responses due to a higher variance in the baseline *OU* process, and a higher variance in resulting threshold-baseline distance in the *eLAT ER* sensory integration process.

To further illustrate the consequences of this result, we simulated the result of a sequential updating scheme, where single parameters of the baseline dynamic were updated according to the expected probability of repetition in a markov process tracking the choice direction outcome. Fig9A depicts this process (left), with the resulting repetition probability approximating a random walk (right), wherein the expected value remains constant, but the variance of the process increases – as such, any given process becomes more likely to have diverged from the expected value as the trial numbers increase, despite the expected value remaining constant. Fig9B-D plots the baseline dynamics (left), anticipatory distributions (middle), and cumulative reaction time distributions (right) for two delay times (750 ms and 1250 ms) for sequential updating of *θ* (Fig9B), *kappa* (Fig9C), and *σ*_*B*_ (Fig9D). Here, two observations should be noted. First, it can be seen that all three parameters can result in changes to the proportion of anticipatory responses, although this may only be apparent if the delay time is sufficiently long to realize this. However, each parameter update has a differential effect on the resulting baseline evidence distributions at different delay times, influencing the trajectory of the evidence accumulation process.

**Figure 9.**
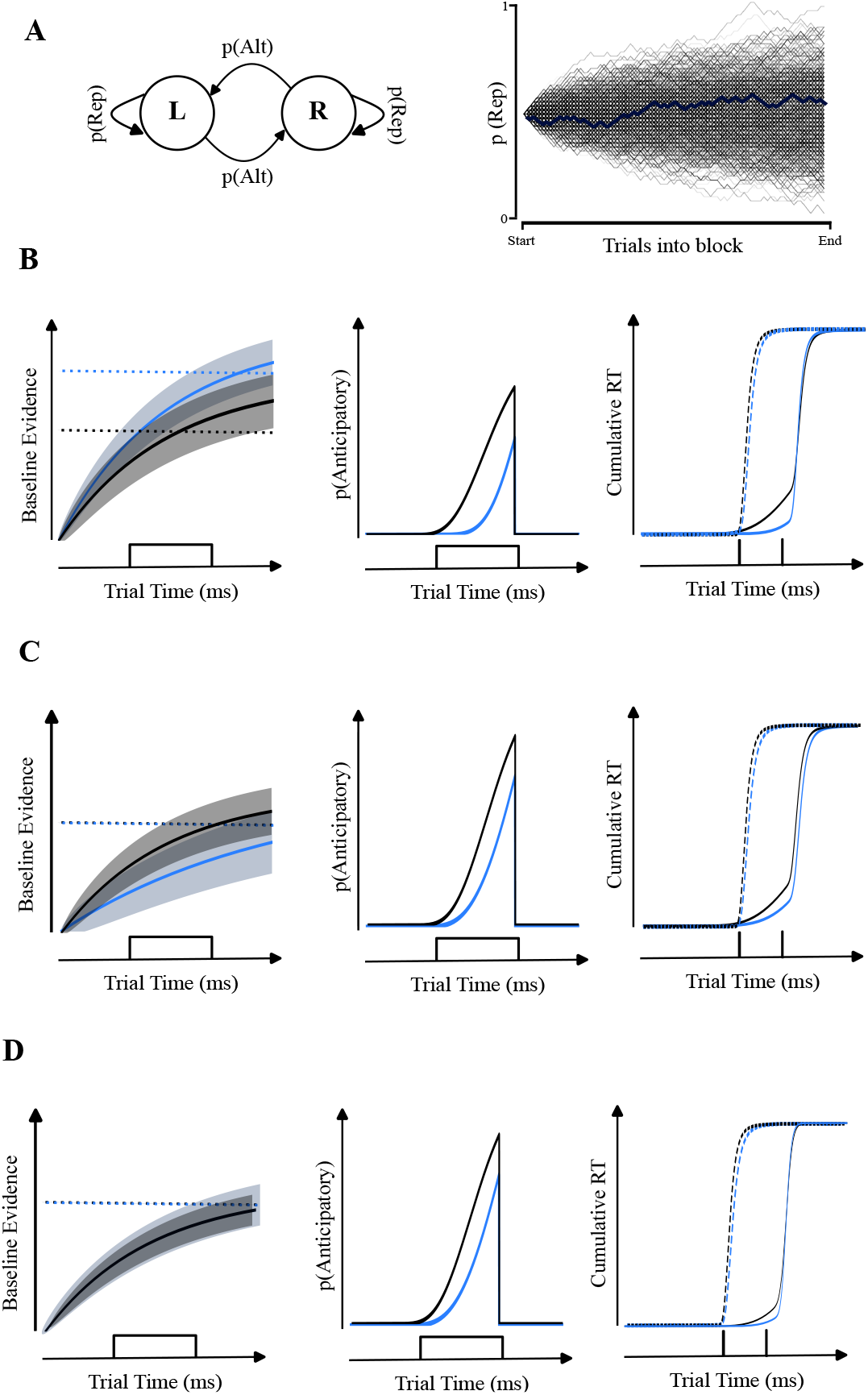
Sequential updating of baseline dynamics parameters. **A:** Depicted is a markov process tracking the probability of choice repetitions and alternations, with the probability of a repetition shown as a random walk over trials (single block in blue, many processes shown in grey). **B** Updating of the mean-reverting parameter *θ*. Black: high repetition probability results in a lower mean-reversion parameter (left panel depicts the baseline dynamics for each), and thus more anticipatory responses (middle panel). The right panel shows reaction time distributions for early target onsets, where the two processes are indistinguishable, and late target onsets, where they are distinguishable. **C:** Updating of the rate parameter *κ*. Higher probability of repetition leads to the baseline dynamics reaching a steady-state faster, thus more anticipatory and faster sensory responses

In light of this, the finding that the variance of the baseline, *σ*_*B*_, bets explained the choice repetition results should be appraised. Like the other parameters, *σ*_*B*_ can result in an increased anticipatory density. However, each parameter has a differential influence on the evidence accumulation process. *theta* influences the *average* baseline, which may or may not be in turn influenced by the delay time, depending on the rate parameter *κ*). In contrast, the variance parameter *σ*_*B*_ influences the variance of the baseline as well as the anticipatory density function. Taken together, this suggests that the choice repetition effects were best explained by a combination of increased anticipatory responses and an increased variance of the thresholdbaseline distance in the extended LATER model, as has been previously reported to underlie pre-target processes such as prediction (Nakahara et al., 2006).

## Discussion

In this paper, we combined behavioural experiments with computational modeling to show that predictions of sensory timing influence how evidence accumulation is gated during saccadic choice, even in a randomized task where history effects may not be expected. In a free choice saccade task to randomized targets presented with variable delays and variable asynchronies (Noudoost and Moore, 2011; Schiller and Chou, 1998; Soltani et al., 2016). We found evidence for an interaction between trial history effects based on the choice sequence, and the influence of the delay period (obesrved via anticipatory responses and delaydependent changes to evidence accumulation). We developed an extended version of a gated accumulator model (Purcell et al., 2012; Schall et al., 2011) to account for this behaviour. In doing so, this work provides a technical advancement on the evidence accumulation framework by explicitly modeling reaction times in reference to a delay interval, rather than as aligned to the onset of a stimulus. This allowed us to estimate latent dynamics in the baseline of an evidence accumulator that may go unobserved in classic accumulator models that align the decision process to the onset of a repeated stimulus. By fitting the model to data, we found that different phenomena in saccadic choice – anticipation, urgency, and history effects – could be accounted for in our data through variance in the baseline dynamics occuring prior to the onset of evidence accumulation. //

Anticipating the timing of sensory information is useful when interacting with recurring patterns in the environment (Nobre and van Ede, 2018), particularly in the presence of delays. In this study, we found that anticipation reflected a balance between lowering the amount of evidence required to trigger a saccade and inhibiting movement generation prior to the appearance of sensory information. The onset of evidence accumulation has been suggested to be under active control and subject to task-specific modulation (Teichert et al., 2016; van den Brink et al., 2021), but in the analysis of choice tasks, anticipatory responses are usually treated as outliers to be discard – as such, the relationship between anticipation and the evidence accumulation may have been underappreciated. When it has, anticipation has been explained as arising from an independent and parallel process (Hernández-Navarro et al., 2021), supporting claims for a mechanistic division between proactive and reactive processing (J. S. B. T. Evans, 2008; Hikosaka and Isoda, 2010). In contrast, our model explicitly links anticipatory processing and evidence accumulation as a dependent process, in accordance with our experimental evidence. However, future work is required to assess if the dependence of baseline evidence and anticipatory response generation is strictly coupled by a common physical process, or only coupled through a joint correlation with another unmeasured factor (Caie and Blohm, 2024).

We also report urgency-like effects in our task, that is a decrease in reaction time as a function of the length of the delay period. We reason this reflected the increasing probability of a target appearing in our task as time goes on. In accumulator frameworks, urgency has classically been associated with a change in the rate of evidence integration (Reddi and Carpenter, 2000)). An additional class of models eschews evidence accumulation altogether, instead arguing that urgency is represented as an independent signal that is combined with sequentially-sampled sensory information (Cisek et al., 2009). Differentiating these models empirically has been a matter of debate (Carland et al., 2015; N. J. Evans et al., 2017; Winkel et al., 2014). As we have not performed a formal model comparison with other potential post-target dynamics, it is unclear to what degree the models can be dissociated empirically through behavioural data alone, particularly without manipulating the strength of sensory information over time (Cisek et al., 2009). Thus, caution is warranted in interpreting the mathematical form of the accumulation process we model as a necessary component of neural processing, rather than a sufficient explanation of the data we present.

The trial-to-trial use of prediction we show evidence for is suboptimal with respect to the instructions and statistics of the task. This sort of sequential dependency is often likened to model-free learning, i.e. updating behaviour purely based on the outcome of the previous choice. The alternative, model-based learning, requires forming an internal representation of the statistics of a task. Model-free strategies have been the *de facto* metaphor for incremental learning in neural systems (Barto, 1995; Montague et al., 1996; Neftci and Averbeck, 2019; Schultz, 1998), while ‘models’ are invoked to explain the learning of more complex structures (Acuña and Schrater, 2010; Daw et al., 2002; Doya, 1999; Harlow, 1949; Nakahara, 2014; Nakahara and Hikosaka, 2012; Tolman, 1948). Model-free strategies, defined this way, are particularly useful for novel or volatile environments, as they can adapt to structures that may not be known *a priori*. For example, a model-free learner in our task could track the recent history of the free choice task with a trial-trial update of the probability of the left target appearing before the right. The disadvantage of such an approach in our task is that while this may be a good approach to learn the *average* statistics of the task, the probability of being biased from the mean on any single trajectory would approximate a random walk, thus it would actually *increases* over time (Lévy, 1940). Thus, a model-free learning algorithm may do a poor job of learning the statistics of the task if given few opportunities to do so, and sequential biases in randomized tasks would emerge as a by-product of the learning process.

Environmental changes occur over multiple timescales (Kording et al., 2007; Murray et al., 2014), so the time horizon with which that change is considered is critical in the ongoing evaluation of behavior (Cheong et al., 2016; Levins, 1968). When making inferences about model-free or model-based learning, it can be convenient to categorize learning by the timescales over which they change: model-free behaviour closely resembles the immediate past, while model-based learning may not. It is worth noting that while model-free learning *is* based on the recent past, learning based on the recent past is not *all* model-free. While modelfree learning refers to an update based solely on the outcome of a previous event, fast-timescale updating could reflect a nuanced model of the volatility of an environment, where rapid switching is favoured in or outside the contexts of a lab task. Although we may be able to quantify the timescale of learning through correlating choices with recent and long-term history, this does not mean we can falsify the generative model (Caie and Blohm, 2024), where what appears to be a suboptimal lingering effect of the previous trial may actually be the output of a learning rule of arbitrary complexity. Indeed, even in simple neural circuits such as those studied in the classic studies of the crab and lobster stomatogastric ganglia, very different physical processes (i.e. synaptic conductances) were shown to be capable of leading to similar emergent circuit properties (**prinz_similar_200**; Marder, 2011) . Such *redundancy* should similarly be considered in future work evaluating history-dependence in gated accumulation – can multiple different across-trial learning schemes describe trial-averaged data equally well? Thus, while we can conclude that our gated accumulation model can describe the trial-averaged statistics of history-dependence we observe, our results do not determine anything about specific learning rules or cost functions that gave rise to the history-dependence, nor *why* participants failed to behave in an algorithmically random fashion.

Our model is statistical in nature, but was inspired by a biophysical model that was originally proposed as a model of the transformation of visual information into accumulated evidence in the frontal eye fields (FEF) (Purcell et al., 2012; Schall et al., 2011). Recording single-neuron activity following the onset of a visual stimuli evokes transient changes to the firing rate of some FEF neurons (Schall et al., 2011; Schmolesky et al., 1998) at latencies between 50 and 100ms. These ‘visual’ neurons (but see Lowe and Schall, 2018) are thought to be located in superficial cortical layers, which receive visual input from intraparietal sulcus (Huerta et al., 1987; Stanton et al., 1995), extrastriate cortex (Jouve et al., 1998; Schall et al., 1995), and the mediodorsal nucleus of the thalamus (Huerta et al., 1986), amongst others. Pyramidal cells in layer V FEF, on the other hand, innervate the superior colliculus (Sommer and Wurtz, 1998) and basal ganglia (Segraves and Goldberg, 1987); neurons of this category are thought to correspond with movement neurons (Ferraina et al., 2002; Segraves and Goldberg, 1987; Sommer and Wurtz, 2000), which begin a ramping discharge after the visual response, reaching a threshold firing rate that correlates with saccade onset (Hanes and Schall, 1996; Hanes et al., 1998). Given the similar profiles of these motor neurons and accumulator models, it has been suggested that the delay in FEF motor neuron activity reflects a transformation of visual information into a motor command. Gated accumulation in FEF was proposed as a mechanism by which ongoing visual transients are filtered from the accumulation of evidence, allowing only salient information to be passed on. The model captures the timecourse of firing rates from visual and motor neurons in different speed-accuracy conditions. The neural implementation of such transformations are a matter of debate (Sajad et al., 2020; Schall et al., 2011).

Our work suggests that the temporal dynamics of this transformation involve a prediction of sensory timing sensitive to trial by trial changes. Classically, the striatum is associated with trial by trial learning (Hwang et al., 2019; Kawagoe et al., 1998; Kravitz et al., 2012; Schultz, 1998; Skelin et al., 2014; Yttri and Dudman, 2016; Yttri and Dudman, 2018). In the striatum, the substantia nigra tonically inhibits collicular activity necessary for saccade generation; this inhibition is released following bursting activity from the caudate, which is reciprocally connected to cortical regions responsible for saccade planning, such as FEF (Hikosaka and Wurtz, 1983). Thus, in addition to reinforcement learning, the striatum is also thought to be important for balancing the facilitation and inhibition of motor plans. Layer V of FEF is reciprocally connected with the striatum (Parthasarathy et al., 1992; Stanton et al., 1988). Pharmacological manipulation of dopamine D2 receptors, primarily found in deep layers, in FEF resulted in a trial-trial repetition bias in the same free choice saccade task used in this paper, while D1 receptors, distributed throughout cortex, resulted in an alternation bias that is typically correlated with sensory processing (Fecteau et al., 2004; Soltani et al., 2013), suggesting that the behavioural dissociation observed in the saccadic system between these biases has a physical correlate.

The striatum is also implicated in timing behaviour. Electrophysiological evidence suggests that monkey striatum amplifies decision dynamics in cortex as a function of urgency (Thura and Cisek, 2017). Timekeeping deficits are also correlated with damage to the striatum in humans, along with classical symptoms of impaired movement initiation (for review see Buhusi and Meck, 2005). So in addition to inhibition of movement and reward-based learning, there is evidence for striatal involvement in temporal processing. Schall and colleagues suggested the striatum as a potential physiological source for providing an inhibitory signal to movement neurons based on the strength of ongoing visual information, along with axo-axonal shunting and intermediary interneurons within FEF (Schall et al., 2011). In our model fitting results, we found that the variance in the baseline dynamics of the gated accumulator best explained the choice repetition data. Given the importance of spiking variability on preparatory eye movement signals in FEF, future work may assess if and how this variability is modulated by trial history.

Our modelling work adds to a growing list of varieties of accumulator models, many of which have very different epistemic intents. Accumulator models in this respect are in a somewhat unique position, in the sense that they are used as both an abstraction of a physical process that makes predictions about mechanisms, and a statistical model that is used to differentiate underlying physical processes across groups or individuals. Perhaps as a result of this, most formulations of accumulator models contain a mixture of theoretical and technical arguments that combine to fit a reaction time distribution. Technical assumptions are usually justified by mathematical convenience rather than logic or theory, yet are necessary to constrain models such that the selective influence of parameters or different model classes (collapsing bounds, baseline variability, etc) can be dissociated by data. Indeed when these models have technical assumptions removed, they are redundant in the sense that they can represent any reaction time distribution even when restricting some degrees of freedom (Jones and Dzhafarov, 2014).

Part of the difficulty in dissociating models and model parameters comes from an additional degree of freedom that must be allowed when one is forced to consider non-stationarity as an additional source of variability. Previously, it has been suggested that baseline evidence in an accumulator framework can be thought of as analogous to a prior belief over the likelihood of one of a limited number of choice alternatives (Carpenter and Williams, 1995; Gold and Shadlen, 2007), motivating the adoption of reinforcement learning approaches to model trial-trial variability in saccadic reaction times (Kim et al., 2017; McDougle and Collins, 2021; Pedersen et al., 2017). We showed in behavioural data that anticipatory and sensory-guided reaction times are jointly influenced by choice history. In our model, we can account for this dependency by allowing the parameters of the baseline dynamic to remain free across fits to choice sequence data. This is a claim for sufficiency of trial-trial variability to capture sequence effects for a specific data-set, however it does not rule out history-dependent variance in sensory integration, as has been reported elsewhere (Steingroever et al., 2021; Urai et al., 2019). Without ground-truth knowledge of a system’s dynamics, it is difficult to estimate the cause of trial-trial variance in model parameters, simply because the hypothesis space over potential models becomes intractable when forced to consider each individual trial as a singularly-observable outcome of a class of models that is inherently stochastic and designed to explain ensembles of trials (Carpenter, 1999).

Because of this, we wish to elaborate on the distinction between our analysis of choice history, and any conclusions that may be drawn with respect to the functional form of underlying history-dependence. What we do report is a dependence on the outcome of previous trials on anticipatory and sensory-guided reaction times for a particular task. From the perspective of the instructions of the task, history effects may be considered to denote some form of suboptimality in the decision process – as the task statistics were randomized, information from the previous trial contained no evidentiary value beyond providing a single sample of the task statistics. Arguments for *why* history effects persist in different tasks are numerous. Some have argued that history effects in randomized experiments (often binary series) reflect a capacity limitation of the system (Baddeley, 1966; Kahneman and Tversky’, 1972; Tune, 1964. Others reframe the question in the context of optimality by considering these history effects as carry-over from rational behaviour in other contexts where statistical regularity is more prevalent, or question if bias is truly present (Ayton et al., 1989; Kareev, 1992; Lopes, 1982). From our data, there is little to be concluded about *why* history effects are present here, simply because the number of sufficient explanations is such that redundancy is likely (Marder, 2011), and falsification is a challenge (Caie and Blohm, 2024). However, we can conclude that saccadic choice was not independent of previous trials, and this history-dependence could be explained by a gated accumulation model driven by a prediction of the timing of future sensory information.

## Acknowledgments

Financial support for this project was provided by the Natural Sciences and Engineering Research Council of Canada (NSERC), the Canada Foundation for Innovation (CFI), and the Deutsche Forschungsgemeinschaft. The first author would like to thank members of the Computational Vision and Kinematics Group and the Theoretical Cognitive Science Group.

## Notes

### Competing Interest Statement

The authors have declared no competing interest.

### Summary of Updates

Paper has been revised to reflect improvements to the model while under review at the Journal of Neurophysiology. This is the updated version that has been submitted to the journal

